# Mechanosensitive PIEZO2 channels shape coronary artery development

**DOI:** 10.1101/2024.07.08.602502

**Authors:** Mireia Pampols-Perez, Carina Fürst, Oscar Sánchez-Carranza, Elena Cano, Sandra Raimundo, Eric L. Lindberg, Martin Taube, Arnd Heuser, Anje Sporbert, Norbert Hübner, Holger Gerhardt, Gary R. Lewin, Annette Hammes

## Abstract

The coronary arteries develop under substantial mechanical loads. However, the role of mechanosensitive ion channels has barely been addressed in this system. Here we demonstrate the expression of the mechanosensitive ion channel PIEZO2 in specific coronary endothelial cell populations during a crucial phase of vascular modeling. *Piezo2* positive coronary endothelial cells display distinct transcriptional profiles and have mechanically activated ionic currents. Strikingly, *Piezo2^-/-^* mouse embryos and mice with human pathogenic variants of *PIEZO2* display coronary vessel malformations and left ventricular hyperplasia. We conclude that an optimal balance of PIEZO2 channel function is indispensable for coronary vessel formation, integrity, and remodeling and likely for proper cardiac function.

## Main

The branching system of the coronary vasculature has a unique structure adapted to the particular physiology of the heart muscle ^1,2^. During its formation in the embryo and throughout adult life, the coronary vasculature is uniquely challenged by mechanical loads from the heartbeat more than 100,000 times a day and from coronary perfusion pressure. The coronary arteries are particularly susceptible to disease and pathologies of these vessels are a main cause of ischemic heart disease, a leading cause of mortality in Western societies ^3–7^. In this context, a better understanding of embryonic coronary artery formation processes holds the potential to recreate and enhance the developmental process for coronary artery regeneration under disease conditions.

Embryonic coronary vasculature formation involves the intricate orchestration of endothelial cell populations, originating from three primary sources: the sinus venosus (SV), the endocardium, and the (pro)epicardium ^7–10^. While the role of chemotaxis in guiding angiogenesis is well described, also for coronary vasculature ^11–13^, there is emerging evidence that mechanical guidance cues, termed durotaxis, play a role in the patterning and morphogenesis of blood vessels ^14–18^. However, there are only few reports on the presence and function of mechanosensitive ion channels in endothelial cells during vasculogenesis or angiogenesis ^19,20^. In particular, PIEZO1 (piezo-type mechanosensitive ion channel component 1), which is widely expressed in many cell types ^21^, is known to play an important role in the initiation of embryonic vessel formation and function ^20,22–24^. PIEZO1-deficient mice exhibit disrupted vascularization and a failure in the maturation of endothelial cells into larger blood vessels ^23,25,26^. The role of its homolog PIEZO2 (piezo-type mechanosensitive ion channel component 2) in angiogenesis during embryonic development has so far not been explored. However, expression of *Piezo2* in endothelial cells of the brain has been suggested ^22,27,28^ and one study reported PIEZO2 to be involved in tumor angiogenesis ^29^. Ever since the discovery of mechanosensitive PIEZO channels ^21^, PIEZO2 has been primarily characterized as a sensory ion channel, present in a variety of sensory neuron types, the function of which is to sense mechanical forces important for touch, interoception and pain ^30–34^. Importantly, *PIEZO2* pathogenic variants are associated with congenital disorders in humans, including joint, craniofacial, brain, and cardiovascular defects ^35–42^. However, the precise function of mechanosensitive PIEZO2 channels in non-sensory cells, including endothelial cells, remains largely unstudied. In particular, a specific function of mechanosensitive ion channels in coronary artery endothelial cells during embryonic development has not been studied. Here we aimed to investigate the role of mechanosensing mediated by PIEZO2 in the developing cardiovascular system. Using a combination of genetic fate mapping, single cell sequencing, physiological analysis, and light-sheet imaging our study uncovers a unique role for PIEZO2 channels in shaping the coronary vasculature during development. Specifically, we demonstrate that dysfunction or loss of PIEZO2 leads to aberrant coronary artery branching, impaired vessel morphology and cardiac hyperplasia.

## Results

### Fate mapping PIEZO2 cells in the embryonic heart

To delineate the trajectory and contribution of *Piezo2* positive cells in the mouse embryonic heart we used a genetic fate mapping approach in which *Piezo2-Cre* ^32^ drives *tdTomato* expression in *Piezo2* expressing cells and their progeny (*Piezo2-tdTomato* mice) (Fig. 1a). Whole-mount three-dimensional confocal imaging revealed specific populations of tdTomato positive (tdTomato+) cells in the E11.5 heart (Fig. 1b). We found tdTomato+ cells in the sinus venosus (SV) and the nascent coronary plexus (CP) and identified them as endothelial cells positive for endomucin and VE-cadherin (Fig. 1b, arrow heads). At E13.5, the tdTomato+ cell population had expanded ^8,10^ to form a nascent coronary endothelial plexus as evidenced by cells staining positive for the pan-endothelial marker platelet/endothelial cell adhesion molecule 1 (PECAM1) (Fig. 1c). Whole-mount confocal microscopy also revealed that tdTomato+ cells were positive for Dachshund homolog 1 (DACH1), a transcription factor specifically expressed in embryonic coronary endothelial cells ^11,43^ (Fig. 1d). Thus, the *Piezo2* expressing cell lineage specifically contributes to the developing coronary vasculature. At E13.5 the coronary plexus finally connects to the ascending aorta and triggers plexus remodeling into a mature vasculature. At E18.5 tdTomato+ cells now clearly formed the neonatal coronary vasculature as revealed by light-sheet microscopy imaging of the entire heart (Fig. 2a and Extended data Movie 1). Optical sections through the whole mount light-sheet images highlighted the connection of the tdTomato+ fate mapped vessels to the ascending aorta, indicating that these vessels are the coronary arteries (Extended data Movie 2). The coronary vessel character was verified on transverse cryosections and we identified tdTomato+ cells as coronary endothelial cells by FABP4 (fatty acid binding protein 4) staining ^44^ (Fig. 2b). Further characterization showed that tdTomato+ cells contribute to both small arteries in the remodeling zone (DACH1-positive) ^11^ (Fig. 2c) and mature arteries, which are positive for SRY-box transcription factor 17 (SOX17) ^45^ (Fig. 2d). The tdTomato+ cells were not co-stained for endomucin (EMCN) in the endocardium, supporting the coronary nature of these endothelial cells ^46^. We observed other non-endothelial cell types with tdTomato labeling, one in the outflow tract with morphological characteristics of vascular smooth muscle cells (VSMCs). These cells were also positive for alpha-smooth muscle actin (αSMA) confirming their identity as vascular smooth muscle cells. Cells making up all four valves of the heart were also tdTomato+ and were surrounded by endothelial cells, indicating their likely identity as valve interstitial cells ^47^ (Extended data Fig. 1). As *Piezo2*-lineage endothelial cells evidently contribute to coronary vasculature formation, we proceeded to analyze both, *Piezo2* expression and the transcriptional profile of *Piezo2* expressing cells.

**Fig. 1.**
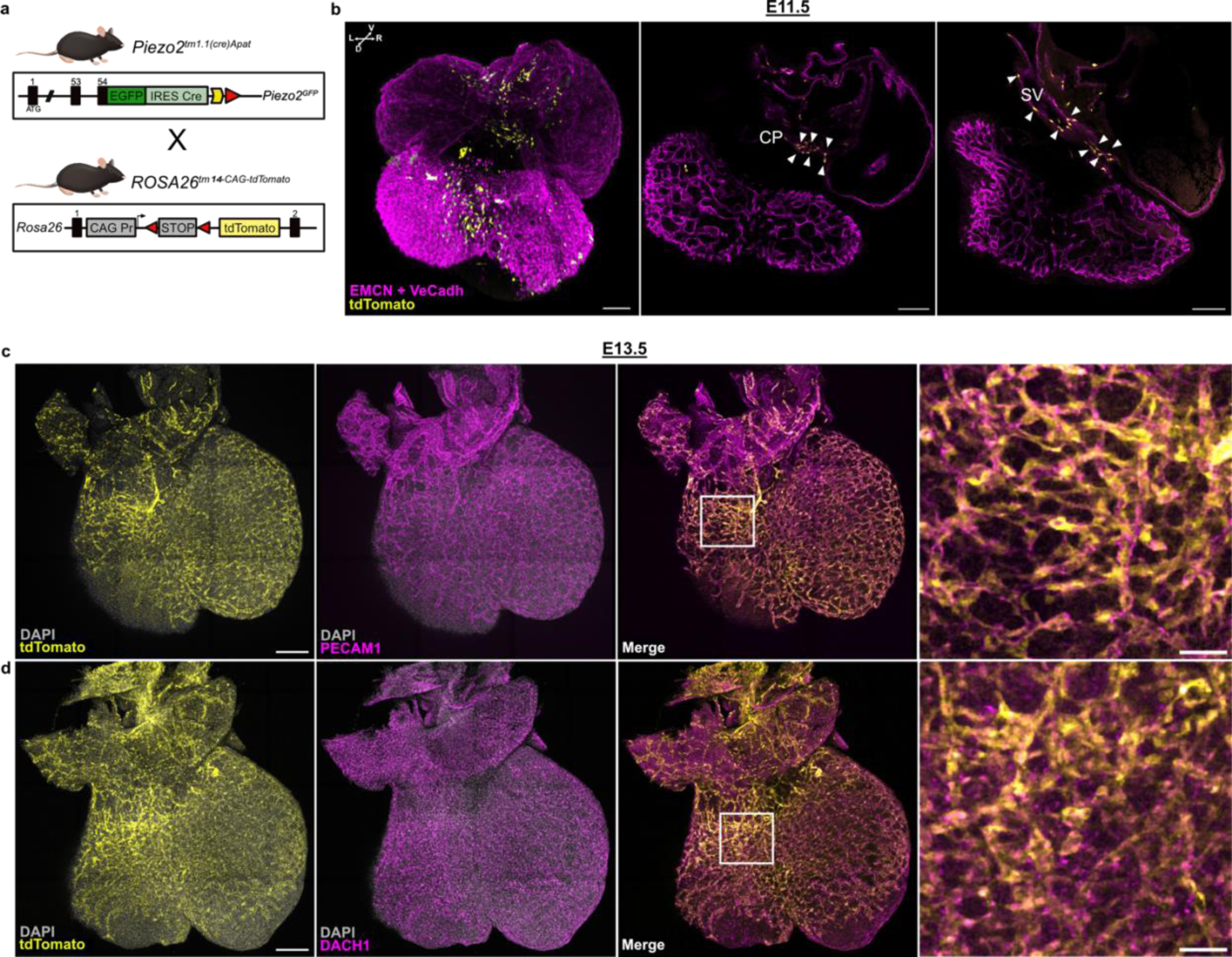
*Piezo2*-driven tdTomato fate mapping in the developing heart. **a**, Outline of the genetic strategy used to generate *Piezo2*-driven tdTomato fate mapping model. **b**, Three-dimensional rendering of representative E11.5 tdTomato+ heart with all cardiac endothelial cells immunolabelled with a cocktail of anti-VeCadherin (VeCadh) and anti-Endomucin (EMCN) antibodies (magenta). tdTomato signal (yellow) was found in the SV and the nascent CP (white arrows in the optical sections) (Scale bar: 100µm) (n = 2). **c**, Three-dimensional rendering of representative E13.5 tdTomato+ heart confirming the endothelial fate (PECAM1 positive, magenta) of tdTomato+ (yellow) cells in the nascent CP (Scale bar: 200µm, 50µm in magnification) (n = 2). **d**, Three-dimensional rendering of a representative E13.5 tdTomato+ heart immunolabelled against the coronary marker DACH1. The co-localization of the tdTomato+ cells (yellow) with DACH1 (magenta) demonstrates that *Piezo2* fate mapped cells were coronary endothelial cells (Scale bar: 200µm, 25µm in magnification) (n=2).

**Fig. 2.**
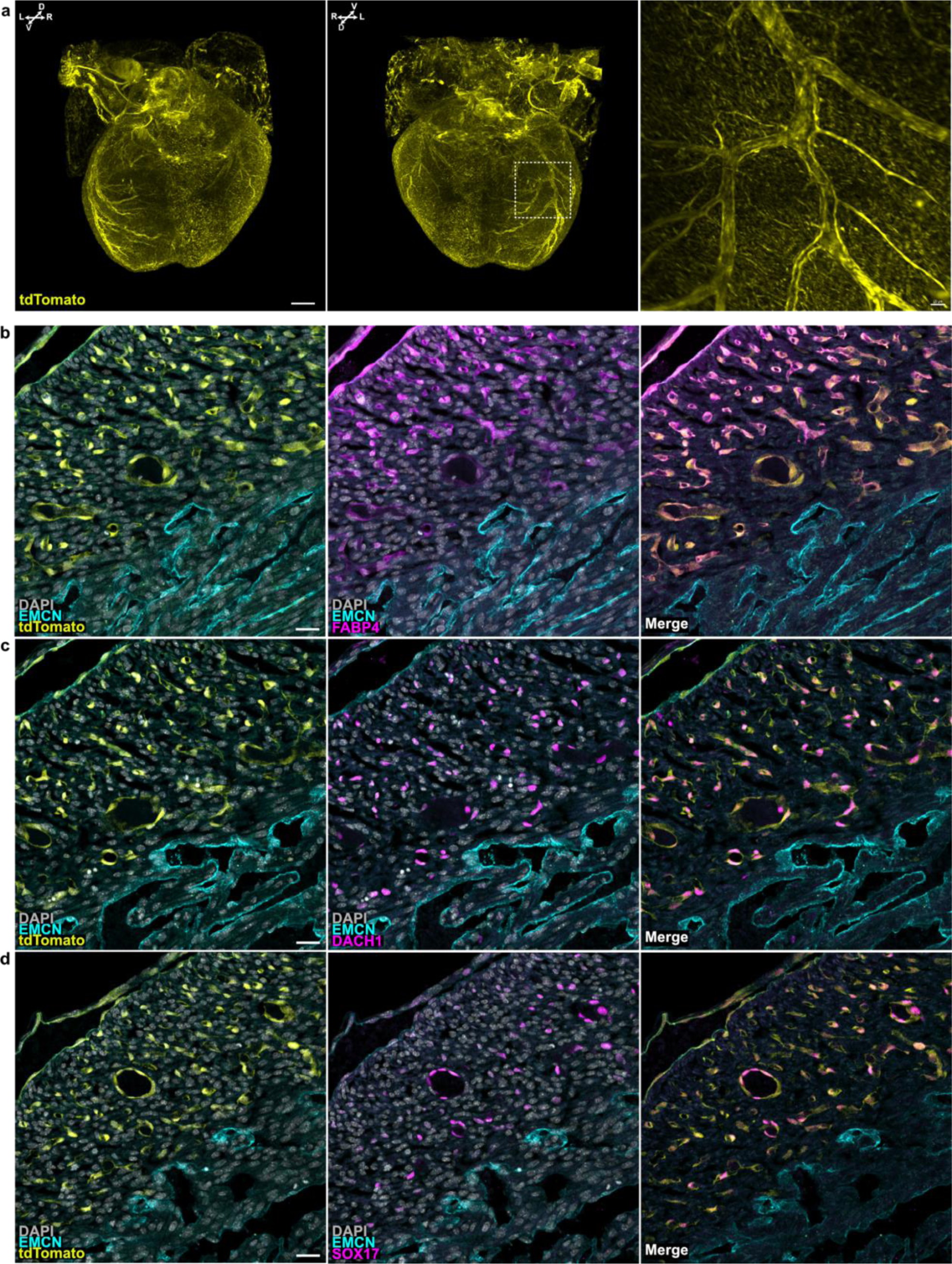
*Piezo2*-driven fate mapping sharply defines the coronary vasculature. **a**, Three-dimensional rendering of a representative E18.5 heart showing that tdTomato+ (yellow) cells contribute to the development of the coronary vasculature (Scale bar: 300µm [3D], 20µm [zoom in] (n = 4). **b - d**, E18.5 tdTomato+ hearts were immunolabelled for endomucin (transversal sections, EMCN, cyan) and three different coronary markers, **b**, FABP4 (magenta). **c**, DACH1 (magenta). **d**, SOX17 (magenta). Immunolabeling revealed that tdTomato+ cells comprise coronary endothelial cells of small (DACH1 positive) and mature (SOX17 positive) arteries, as well as endothelial cells in the coronary FABP4-positive population. tdTomato signals did not co-localize with EMCN in the endocardium (cyan), confirming the coronary endothelial cell nature of the tdTomato+ cells (Scale bars: 25µm) (n = 2).

### Single cell sequencing reveals *Piezo2* expression in specific cell types

While fate mapping provided insights into the trajectories of *Piezo2* expressing cells, examining the transcriptional profile of *Piezo2* expressing cells could help identify the developmental stages in which this mechanosensitive ion channel shapes coronary development. Using single molecule fluorescent in situ hybridization (smFISH) we could show continued *Piezo2* mRNA expression in coronary endothelial cells at E13.5 along with *Dach1* expression (Extended data Fig. 2a). Additionally, single cell RNA datasets (scRNA) from FACS-sorted cardiac endothelial fractions (CD31+ alias PECAM1+/CD45-alias PTPRC^-^) from four different stages: E12.5, E15.5, postnatal day (P)2 and adult (8 weeks old) were used to monitor ongoing *Piezo2* expression ^48^. Unsupervised clustering was performed using the Uniform Manifold Approximation and Projection (UMAP) method and cell type clusters were classified based on the expression profiles of the top differentially expressed genes ^48^ (Fig. 3a). At E12.5 *Piezo2* expression was mainly found in the coronary endothelial cells (CoEC). At E15.5 *Piezo2* expression was mainly found in two sub-clusters of coronary endothelial cells designated CoEC_I and CoEC_II, which are capillary cells ^48^. At postnatal day 2, *Piezo2* expression was still present in the same capillary cell types (CoEC_I and CoEC_II) and in the proliferating cell cluster however, in adult hearts, no significant *Piezo2* expression was found in any of the cell types analyzed (Fig. 3b). These data demonstrate a dynamic *Piezo2* expression pattern in distinctive coronary endothelial cell populations during embryonic heart development.

**Fig. 3.**
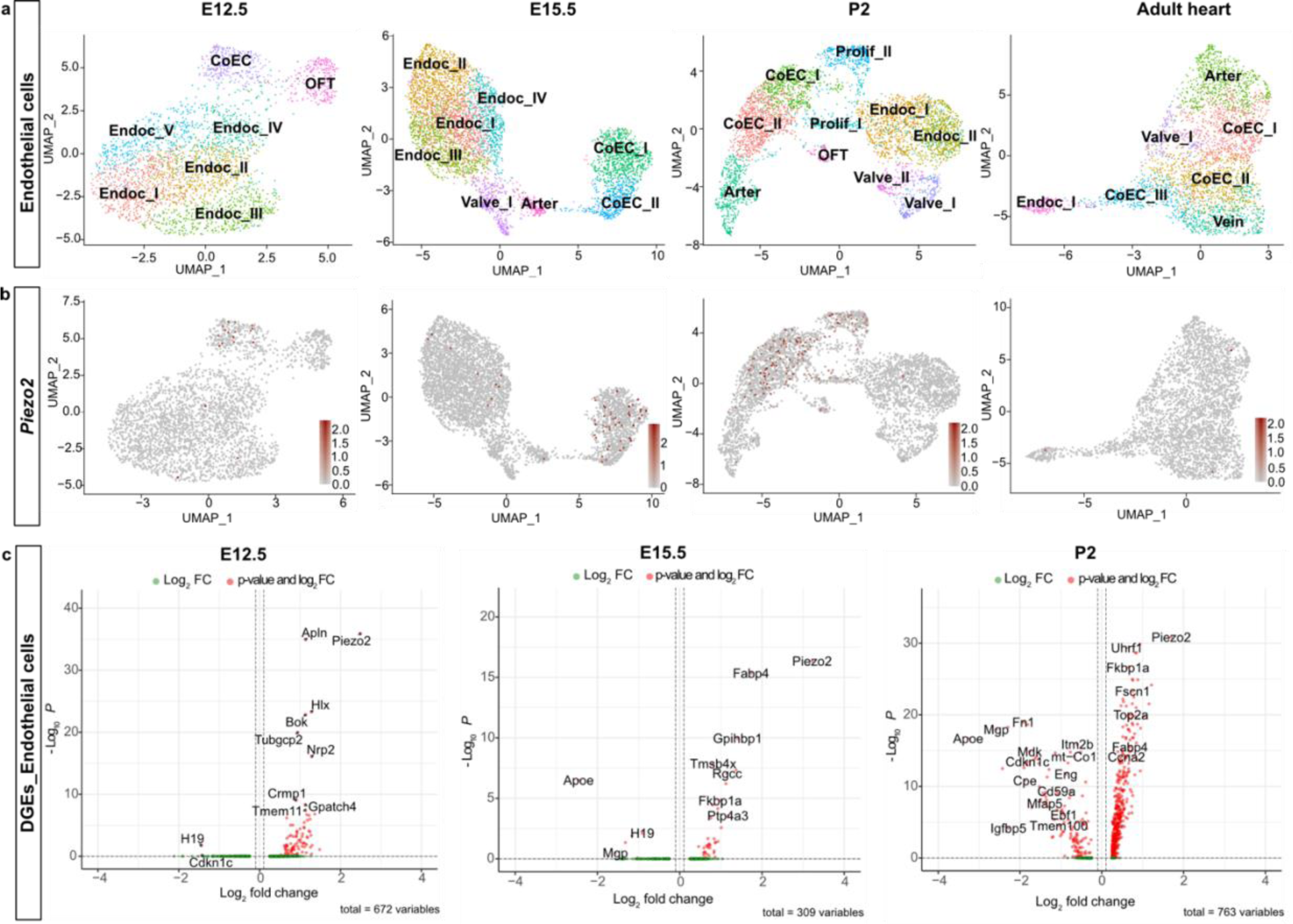
scRNAseq confirmed ongoing expression of *Piezo2* in the developing coronary endothelial cells. **a**, UMAP of cardiac endothelial cells from E12, E15, P2 and murine adult hearts (8 weeks old) showing the following clusters: endocardium (Endoc_I to Endoc_V), coronary endothelium (CoEC_I to CoEC III), proliferating cells (Prolif_I to Prolif_II), valvular endocardium (Valve_I and Valve_II), arterial cells (Arter), venous cells (Vein), and outflow tract cells (OFT). **b**, Representation of *Piezo2* expression in various clusters of cardiac endothelial cells. **c**, Volcano plots showing differentially expressed genes (DEGs) between individual *Piezo2* expressing and *Piezo2* non-expressing endothelial cells.

### *Piezo2* positive coronary artery cells have distinct transcriptional profiles

We next asked if *Piezo2* expressing coronary endothelial cells show a distinct transcriptional profile when compared to coronary endothelial cells lacking *Piezo2* expression (Fig. 3c, Extended data Tables 1 - 3). At E12.5 *Piezo2* positive coronary endothelial cells showed higher expression of genes like *Neuropilin 2* (*Nrp2*), a receptor for semaphorin family members, and *Apelin* (*Apln*), a sprouting marker, that both have been implicated in blood vessel formation ^13,49–53^. We also noticed enrichment of collapsin response mediator protein 1 (*Crmp1*), a protein involved in neuronal migration and development ^54^. *Piezo2* was still enriched at E15.5 along with other well-known markers of coronary endothelial cells like *Fabp4* ^52,55^ and the capillary markers *Rgcc* (regulator of cell cycle) and *Gpihbp1* (glycosylphosphatidylinositol anchored high density lipoprotein binding protein 1) ^56,57^. At P2, *Piezo2* positive cells still showed enriched expression of *Fabp4* and interestingly were positive for fascin actin-bundling protein 1 (*Fscn1*), an actin-bundling protein that can induce membrane protrusions and is involved in cell migration and motility ^58^ (Fig. 3c). The PIEZO2 homolog PIEZO1 is a stretch activated ion channel that is widely expressed in the cardiovascular system ^23,59–61^. Consistently, our analysis revealed that numerous cell types exhibit significant *Piezo1* expression across all stages investigated (Extended data Fig. 2b). In contrast, *Piezo2* expression appeared to be maintained primarily in coronary endothelial cells specifically during the angiogenic phase of heart vascularization. The distinct transcriptional signature of *Piezo2* expressing coronary endothelial cells indicates potential roles in cell migration and motility. These findings provide insights into the molecular mechanisms underlying the development and maturation of coronary endothelial cells, highlighting the importance of PIEZO2 and associated markers in cardiovascular biology.

### Endothelial cells show PIEZO2-like mechanosensitive currents

The function of PIEZO2 in non-sensory endothelial cells has not been studied. PIEZO2 ion channels are very efficiently activated by substrate deflection, but unlike PIEZO1 channels, are poorly activated by membrane stretch ^62–65^. We identified the yolk sac from E13.5 to E18.5 embryos as being rich in tdTomato+ endothelial cells, which may express functional PIEZO2 channels (Extended data Fig. 3). We asked whether these cells possess functional mechanosensitive currents characteristic of PIEZO2 channels. We made whole cell patch clamp recordings from tdTomato+ and tdTomato-yolk sac endothelial cells cultured on pillar arrays to study currents gated by substrate deflection (Fig. 4a) ^62,63^. We also tried cell indentation, a method often used to activate mechanically activated channels ^62,63,66,67^, but the endothelial cells were too small and flat to make this method feasible. Nanometer scale pillar displacements move the substrate-cell contact and are limited to the pillar area (10.3µm^2^). This mechanical stimulus efficiently evoked inward currents with short latencies (< 5 ms) in almost all cells studied, due to the opening of mechanosensitive ion channels sub-adjacent to the stimulated pillus (tdTomato+ 12/13 cells, tdTomato-11/12 cells). Mechanosensitive currents could be classified into three groups with different inactivation time constants similar to what has been shown in sensory neurons ^63^, rapidly adapting (RA) currents (<5 ms), intermediately adapting (IA) currents (5-50 ms), and slowly adapting (SA) currents (>51 ms) (Fig. 4b). Notably, the RA-currents possessed ultra-fast kinetics in that many of the currents inactivated within a millisecond (Fig. 4c). Very fast inactivation is a characteristic property of PIEZO2 channels as in contrast PIEZO1 channel inactivation time constants, following mechanical stimuli, are typically considerably longer ^21,62,63^. Consistently, RA-currents with very fast inactivation kinetics were found to be significantly more frequent in tdTomato+ cells (Chi-squared P<0.05) (Fig 4B). In addition, mean inactivation time constants for all measured currents were faster in tdTomato+ compared to tdTomato-cells (Fig. 4c). The frequency with which mechanosensitive currents could be evoked was higher in tdTomato+ compared to tdTomato-cells indicating enhanced mechanosensitivity in *Piezo2* positive lineage cells. However, no statistically significant differences were observed when comparing deflection-current amplitude relationships (Fig. 4d, Extended data Fig. 4a). In summary, almost all yolk sac endothelial cells possess very sensitive mechanically activated currents, but cells from the *Piezo2* positive lineage possessed more currents with kinetics characteristic of recombinantly expressed PIEZO2 channels ^21,63^. To get closer to the role of mechanosensitive currents in the development of the coronary circulation we managed to isolate tdTomato+ cardiac endothelial cells (cECs) from E13.5 hearts and cultured these cells on pillar arrays. Almost all cells identified as tdTomato+ two weeks after plating exhibited very robust and large mechanically activated currents to substrate deflection (14/15 cells tdTomato+, 7/8 tdTomato-) (Fig. 4a, e - g). However, unlike yolk sac cells, the kinetics and current amplitudes of mechanosensitive currents did not significantly differ between tdTomato+ and tdTomato- cells (Fig. 4e - f). We measured absolute deflection thresholds and currents were evoked by movements of less than 50 nm (Extended data Fig. 4b), a similar sensitivity to that is seen for PIEZO2-dependent touch receptors ^63,68,69^. However, the frequency with which mechanosensitive currents could be evoked was higher in cardiac endothelial cells that were tdTomato+ (Fig. 4g). Thus, there appears to be clear PIEZO2-dependent endothelial cell mechanosensitivity, but other mechanosensitive ion channels like PIEZO1 and ELKIN1 are probably also present ^23,24,70^. Consistent with this idea, we could also measure clear stretch activated currents, typical for PIEZO1 channels, in excised outside-out patches from tdTomato+ yolk sac endothelial cells (Extended data Fig. 4 c - d). Moreover, *Elkin1* transcripts were detected in the same endothelial cell populations in which we had detected *Piezo2* (Extended data Fig. 5).

**Fig. 4.**
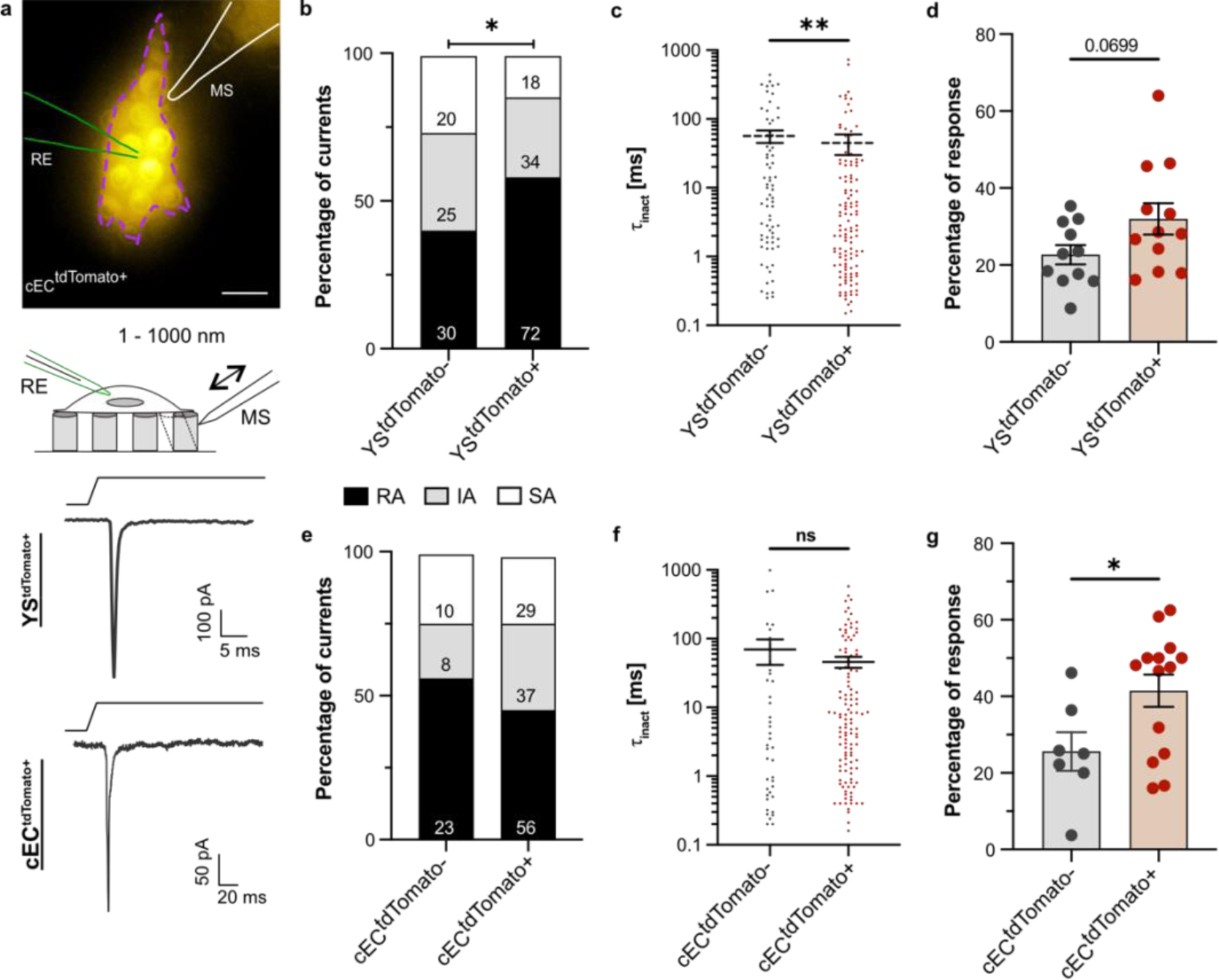
Yolk sac (YS) and cardiac endothelial (cEC) cells displayed endogenous deflection-gated currents. **a**, *above*: Representative picture of a cEC^tdTomato+^ cell cultured on the elastomeric pillar arrays. (RE recording electrode, MS = mechanical stimulator. Scale bar: 10 μm). *below*: Example traces of rapidly adapting (RA) currents from YS^tdTomato+^ and cEC^tdTomato+^ cells recorded with the pillar arrays method. **b**, Stacked histograms show that the proportion of RA currents in YS^tdTomato+^ cells (n = 12) was higher compared to YS^tdTomato-^ cells (n = 11 Chi-squared test, p = 0.02). **c**, YS^tdTomato+^ cells showed deflection-gated currents with faster inactivation kinetics compared to non-red cells (Mann-Whitney test, p = 0.004). **d**, YS^tdTomato+^ cells responded statistically similar to YS^tdTomato-^cells, with a trend that shows that they are slightly more sensitive to mechanical stimulation (t-test, p = 0.0699). **e - f**, cEC^tdTomato+^ (n = 14) and cEC^tdTomato-^ cells (n = 7) did not show differences in the proportion of mechanosensitive currents nor changes in inactivation kinetics. **g**, Box plots showing that cEC^tdTomato+^ cells are more responsive to deflection stimuli compared to cEC^tdTomato-^ cells (Mann-Whitney test, p = 0.0392). (SA = slowly adapting, IA = intermediately adapting).

### Pathogenic *Piezo2* mutations causes cardiac hyperplasia

We next asked whether PIEZO2 channels are necessary for normal heart development. Since *Piezo2^-/-^* mutant mice die perinatally ^30,32,71^, we examined hearts from these mice at E18.5. *Piezo2^-/-^* mutant hearts were smaller and shorter at birth compared to wild-type (WT) *Piezo2^+/+^* littermates (Fig. 5a - b). However, measuring heart/body weight ratios, we found ratios were increased in mutants, indicating that *Piezo2^-/-^* embryonic hearts were heavier than WT hearts (Fig. 5c). We made a detailed morphological analysis using the endocardial marker endomucin (EMCN) to differentiate between the trabecular myocardium (TM) and compact myocardium (CM) (Fig. 5d). The results revealed that *Piezo2^-/-^* hearts showed clear ventricular hyperplasia which was most evident for the left ventricle (LV) (Fig. 5e - f). In addition, the intraventricular septum (IVS) was also significantly thickened (Fig. 5g). No obvious outflow tract defects, such as common arterial trunk and double outflow tract right ventricle were observed in the *Piezo2^-/-^* mutants. Pulmonary artery and ascending aorta diameter were comparable between WT and *Piezo2^-/-^* hearts (Extended data Fig. 6). Longitudinal and transverse sections of the valves showed clear separation of the leaflets (Extended data Fig. 6).

**Fig. 5.**
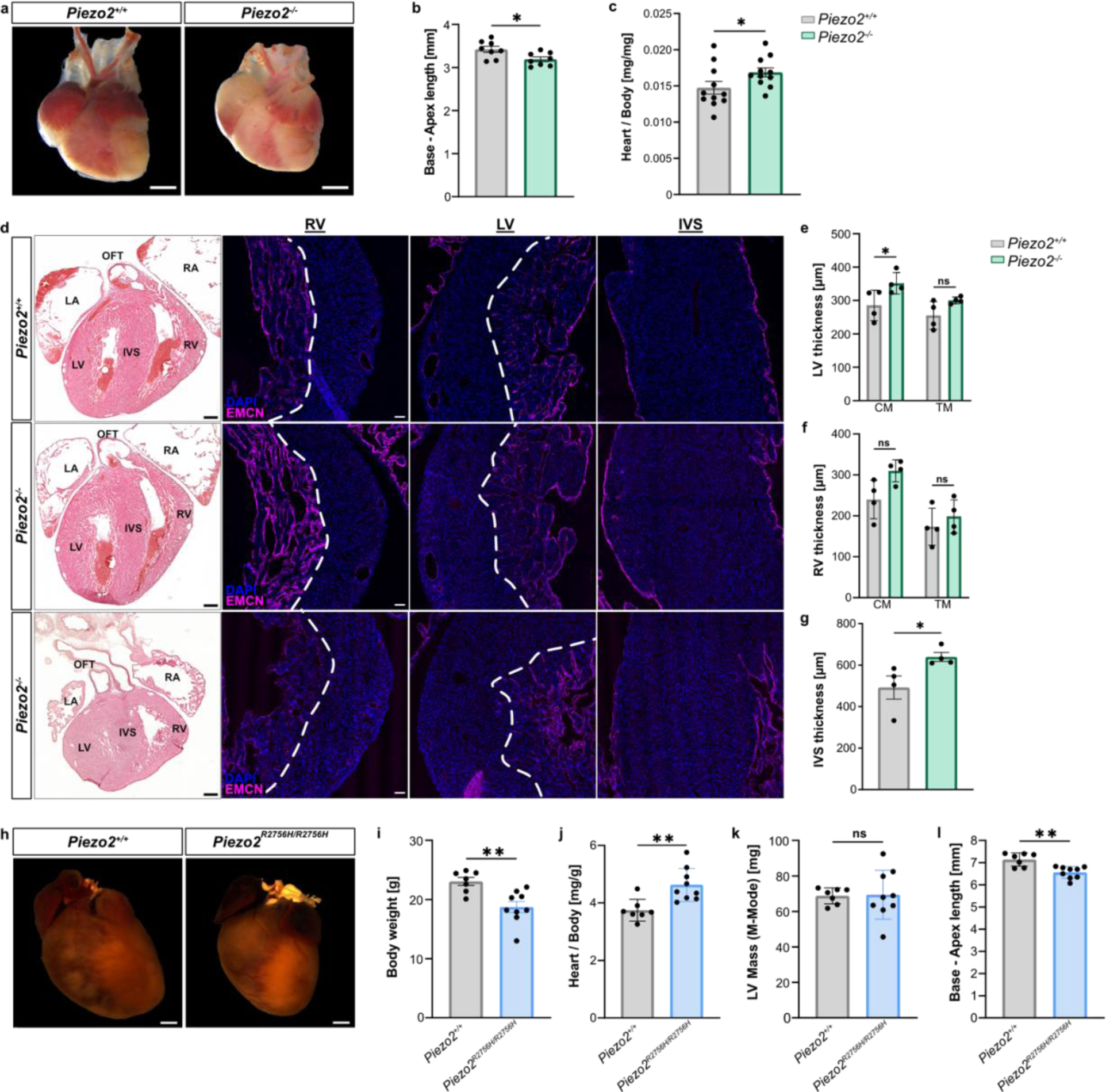
*Piezo2* mutations cause cardiac hyperplasia. **a**, E18.5 *Piezo2^+/+^* and *Piezo2^-/-^* macroscopic analysis. **b**, *Piezo2^-/-^* mutant hearts display a shorter base to apex length (n = 8 per genotype, Mann-Whitney test, p = 0.02). **c**, *Piezo2^-/-^* mutant mice had increased heart-to-body weight ratio (n = 11 per genotype, Mann-Whitney test, p = 0.04). **d**, Ventricular wall thickness was measured from E18.5 *Piezo2*^+/+^ and *Piezo2*^-/-^ hearts immunolabelled against EMCN (magenta) to distinguish between compact and trabecular myocardium (CM and TM, respectively) (Scale bar: 50µm). Hematoxylin and eosin staining was performed for an overall morphological heart assessment (Scale bar: 250µm). Two representative *Piezo2*^-/-^ hearts are shown with a milder hyperplasia phenotype (middle row) and a severe hyperplasia phenotype (last row), including the presence of an aberrant outflow tract (OFT). **e** - **g**, *Piezo2*^+/+^ and *Piezo2*^-/-^ (n = 4 per genotype) right ventricle (RV), left ventricle (LV) and interventricular septum (IVS) wall thickness quantification show that E18.5 *Piezo2*^-/-^ present a thicker LV wall and IVS, indicating cardiac hyperplasia (LV, two-way ANOVA, p = 0.04 (CM) and p = 0.17 (TM); RV, two-way ANOVA, p = 0.06 (CM) and p = 0.64 (TM); IVS, Mann-Whitney test, p = 0.02). **h**, Representative macroscopic heart images of 20-week-old mice carrying a *Piezo2* knock-in gain-of-function mutation (p. Arg2756His) and *Piezo2^+/+^* mice (Scale bar: 1mm). **i** - **l**, Echocardiography of 10 weeks old *Piezo2^+/+^* and *Piezo2^R2756H/R2756H^* mice. **i**, *Piezo2^R2756H/R2756H^* mice’s body weight is significantly less than *Piezo2*^+/+^ mice (Mann-Whitney test, p = 0.003). **j,** The heart-to-body weight ratio is increased in *Piezo2^R2756H/R2756H^* mutants compared to *Piezo2*^+/+^ (Mann-Whitney test, p = 0.003). **k**, Both groups had similar LV mass (Mann-Whitney test, p = 0.83). **l**, Heart size was measured from base to apex showing that *Piezo2^R2756H/R2756H^* hearts are smaller compared to *Piezo2^+/+^* (Mann-Whitney test, p = 0.004), indicating hyperplasia (n = 7 *Piezo2^+/+^* and n = 9 *Piezo2^R2756H/R2756H^*).

**Fig. 6.**
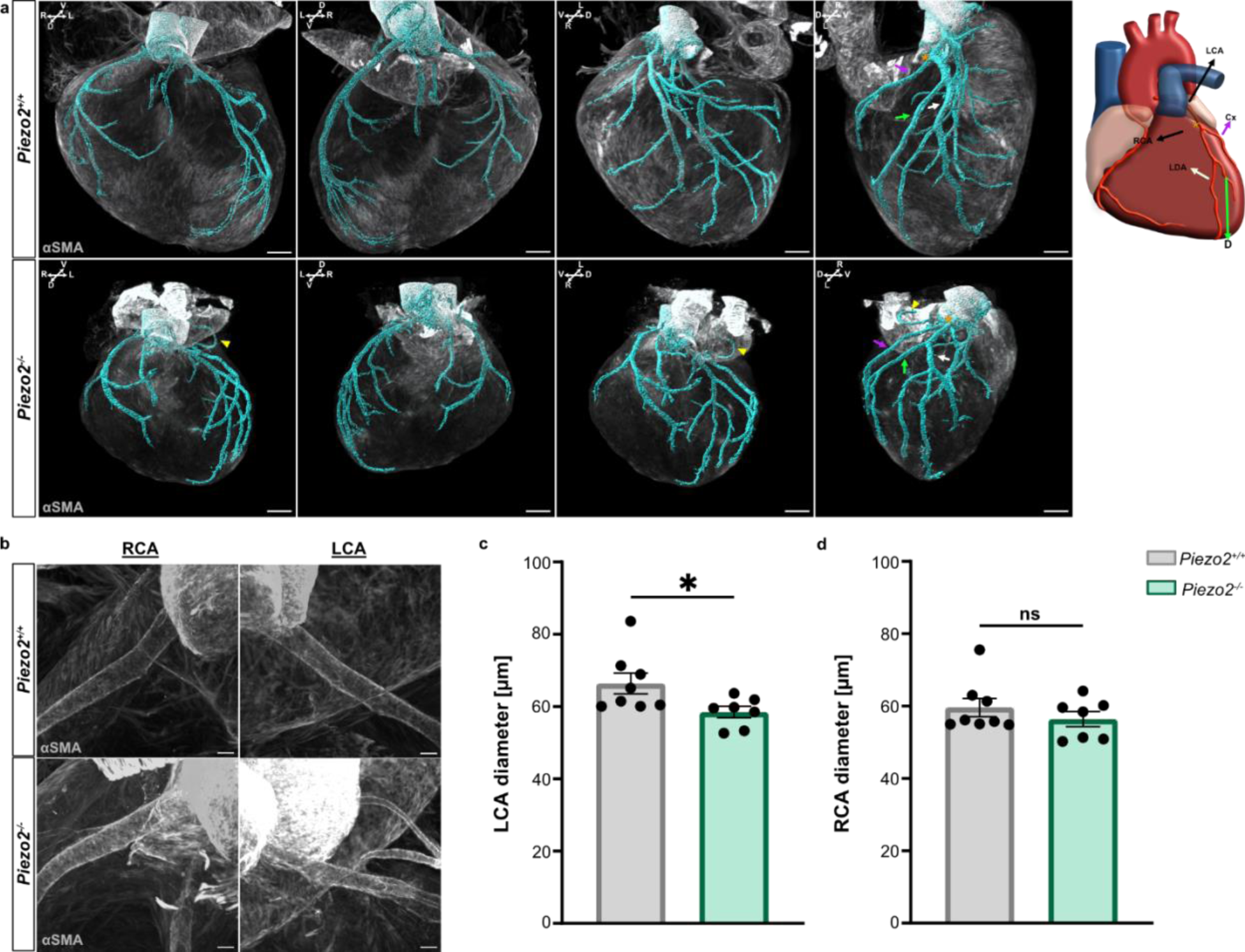
*Piezo2* loss of function mutations alter coronary artery development. **a**, Three-dimensional rendering of representative *Piezo2^+/+^* and *Piezo2^-/-^* E18.5 hearts immunolabelled against αSMA (gray), a smooth muscle marker, used for coronary vasculature segmentation (cyan) (Scale bar: 250µm). The schematic shows the right and left coronary arteries (RCA and LCA, respectively) and the LCA main branches: the circumflex artery (Cx, magenta arrow), the diagonal artery (D, green arrow) and the left descending artery (LDA, white arrow). The orange asterisk depicts the Cx bifurcation point. The yellow arrowhead in the mutant heart points to an aberrant ectopic coronary branch in a *Piezo2^-/-^* heart. **b** - **d**, RCA and LCA diameters were measured and compared between *Piezo2^+/+^* (n = 8) and *Piezo2^-/-^* (n = 7). *Piezo2^-/-^* present with a significantly smaller left LCA diameter (Mann-Whitney test, p = 0.02 [LCA] and p = 0.53 [RCA]). Representative images are shown in **b** (Scale bar: 50µm).

Rare *PIEZO2* gain-of-function pathogenic variants in humans are associated with neurodevelopmental disorders such as Gordon syndrome and Marden-Walker syndrome as well as Distal Arthrogryposis ^35,36,38–40,72^. We generated mice carrying a pathogenic homozygous gain-of-function mutation (*Piezo2^R2756H/R2756H^*) that dramatically sensitizes the activation of endogenous PIEZO2 channels by relieving voltage block of the channel ^73^. Homozygous *Piezo2^R2756H/R2756H^* mutants are viable and so we examined the hearts of these animals at 20 weeks of age post-mortem (Fig. 5h). Similarly, to embryonic *Piezo2^-/-^* hearts, we found that the *Piezo2^R2756H/R2756H^* mutant hearts were smaller and shorter than hearts from age-matched WT littermates. We noted that *Piezo2^R2756H/R2756H^* mutant mice at 10 weeks weighed significantly less than WT littermates (Fig 5i). Indeed, echocardiography measurements from living *Piezo2^R2756H/R2756H^* and WT littermate mice at 10 weeks of age showed that *Piezo2^R2756H/R2756H^* mutant mice displayed a significantly elevated heart to body weight ratio compared to controls (Fig. 5i - j). LV mass was comparable between genotypes (Fig. 5k), but mutant hearts were significantly shorter than WT hearts (Fig. 5l). These results suggest that both loss-and gain-of-function of PIEZO2 can lead to macroscopic hyperplastic changes in the embryonic and adult heart.

### PIEZO2 shapes normal coronary artery development

The LV hyperplasia observed in *Piezo2^-/-^* mice, is a phenomenon that can result from reduced perfusion of the cardiac muscle. Since PIEZO2 is highly specific to the coronary endothelium, we hypothesized that PIEZO2 mechanosensory function in these cells guides the normal assembly of the coronary vasculature. We performed whole-organ immunostaining against αSMA in iDISCO cleared embryonic hearts. Using light-sheet imaging and vessel segmentation we could visualize the entire coronary vasculature in WT and *Piezo2^-/-^* hearts (Fig. 6a - b, Extended data Fig. 7, Extended data Movies 3 - 8). The majority of the main branches of the coronary artery network follow a stereotypical pattern ^74,75^. From the coronary artery ostium at the left coronary cusp of the ascending aorta, the left main coronary artery (LCA) extends ventrally before the first bifurcation into the left anterior descending artery (LDA) and left circumflex artery (Cx). Further ventrally, the diagonal branch (D) bifurcates from the LDA. Conversely, the right coronary artery (RCA) connects to the aorta at the right coronary cusp.

In *Piezo2^-/-^* hearts the coronary vasculature topology was clearly altered compared to WT. In particular, in *Piezo2^-/-^* hearts, the branching of the left coronary artery into the circumflex artery (Cx), the diagonal branch (D), and the left anterior descending artery (LDA) did not follow the pattern seen in WT. There was no clear distinction between the main LDA and its branches in the mutant heart compared to the WT (Fig. 6a, white arrow, Extended data Fig. 6d, white arrow). The branchpoints in mutant vessels resembled trifurcations rather than bifurcations. Moreover, the left circumflex artery branched off from the main coronary artery further to the anterior pole (Fig. 6a, orange asterisk; Extended data Fig. 6, orange asterisk). Additionally, in mutant hearts, an ectopic artery branching off the Cx projecting towards the anterior pole of the heart was observed (Fig. 6a, yellow arrowhead).

Most importantly, the light-sheet reconstruction indicated that *Piezo2^-/-^* hearts have more constricted coronary vessels. Since LV hyperplasia was most prominent in *Piezo2^-/-^* hearts we quantified the average diameter of the left and right main coronary artery over 600µm from the ostia (Fig. 6c - d). We observed significant narrowing of the left coronary artery diameter in *Piezo2^-/-^* hearts compared to WT. Taken together we conclude that the mechanosensitive ion channel PIEZO2 is necessary for normal assembly of the cardiac endothelium to form an adequate vasculature supply to the developing myocardium.

## Discussion

Our investigation reveals a critical role of mechanosensitive PIEZO2 function in the formation of the coronary vasculature, particularly in developing coronary endothelial cells. We show specific expression of *Piezo2* in coronary endothelial cells, which exhibit unique expression profiles characterized by the enrichment of genes involved in cell migration, guidance, and angiogenesis, including *neuropilin 2*, *apelin*, and *Crmp1* ^50,53,54,76^. Neuropilin 2, originally recognized for its role in neuronal guidance ^53^, also influences angiogenesis and likely guides endothelial cell migration and vessel formation ^50^. Similarly, apelin, a hormone known for supporting angiogenesis, enhances endothelial cell survival and stabilizes blood vessels ^76–78^. Additionally, CRMP isoforms, initially associated with neuronal development ^54,79^, are present in endothelial cells, where they likely contribute to cytoskeletal regulation, cell movement, and angiogenesis. Of note, *PIEZO2* expression has been reported in human cultured coronary artery endothelial cells ^80^ and in endothelial cells of cardiac tissue samples from transplanted human hearts ^81^.

In a scRNA-seq analysis of P7 murine brain endothelial cells, it was shown that *Piezo2* was amongst the 25 most enriched tip cell transcripts ^82^. PIEZO2 has been implicated in transducing physical cues into mechanobiological responses, such as cytoskeletal rearrangements ^83,84^. It is therefore plausible to hypothesize that PIEZO2 is needed to guide endothelial navigation at the angiogenic front of the coronary vascular plexus in response to mechanical cues.

Chemical signaling (chemotaxis), particularly through pathways like VEGFA-VEGFR2, BMP2, and CXCL12/CXCR4, is well-established in orchestrating endothelial cell movements underlying angiogenesis in various organs, including the heart ^11–13,46^. However, there is increasing evidence that combinatorial effects of chemotaxis and ion channel-mediated durotaxis fine-tune cellular migration and navigation in diverse cellular systems ^17,85–87^. In particular, the study by Canales Coutiño and Mayor in *Xenopus* showed that Piezo1 cooperates with chemotactic signaling cues to control neural crest migration ^85^. PIEZO2 was also shown to balance mechanosensitivity and chemosensitivity at the blood-tumor barrier in medulloblastoma ^88^.

Indeed, endothelial PIEZO2 could be a major sensor of mechanical cues in the developing myocardium, facilitating cellular migration and motility through durotaxis. We propose that mechanical cues sensed by *Piezo2* positive coronary endothelial cells are crucial for precise cellular navigation during coronary vasculature plexus formation and remodeling.

Our functional analysis demonstrated that *Piezo2* positive endothelial cells possess fast mechanosensitive currents that have sensitivities and kinetics similar to *Piezo2*-dependent currents in sensory cells ^63,68,73^. However, many cardiac endothelial cells that had never expressed *Piezo2* also exhibited mechanosensitive currents (Fig. 4), showing that PIEZO2 is only one of several functional mechanosensitive channels in cardiac endothelial cells ^20^. Indeed, our data show that *Elkin1* and *Piezo1* are also expressed in cardiac endothelial cells, and both channels are efficiently gated by substrate deflection or membrane stretch ^63,70^. Nevertheless, it is striking that PIEZO2 loss-and gain-of-function experiments revealed a non-redundant role of endothelial PIEZO2-dependent mechanosensitivity in shaping coronary artery formation and branching.

We observed aberrant coronary artery branching and a decreased left coronary artery diameter in the absence of *Piezo2* expression. We propose that cardiac muscle perfusion deficits result from coronary anomalies in *Piezo2^-/-^* mutants and as a consequence the mice show heart malformations and cardiac hyperplasia (Fig. 5 a - g). Human pathogenic gain-of-function mutations in *PIEZO2,* documented in human patients ^35,39,72^, dramatically increase the channels’ sensitivity to mechanical force ^73^. By introducing this gain of function mutation into mice we show here that PIEZO2 hypersensitivity can also lead to a disruption of heart development (Fig. 5 h - l).

While the main causes of coronary artery disease are adult onset conditions such as atherosclerosis, hypertension and diabetes due to lifestyle modifications, genetic predisposition can also play an important role in disease etiology ^89–92^. Congenital coronary artery defects have been implicated in cardiac hyperplasia/hypertrophy ^93–95^ and although they might be mild without any symptoms during development and in adolescence they are considered important factors that can predispose individuals to cardiac disease in adult life ^96^. Indeed the heart malformations that we have observed are likely to contribute to the early perinatally mortality in *Piezo2^-/-^* mice which was attributed to failure to inflate the lungs after birth ^30,32,71^.

The importance of PIEZO2 in a variety of sensory functions, including pain, makes it a potentially attractive target for drug therapies ^97^. However, our findings indicate that PIEZO2 is also crucial for the development of coronary arteries in the heart. Considering that developmental programs are often reused in regeneration and remodeling conditions, PIEZO2 might be a promising candidate to be re-expressed in the adult heart to facilitate angiogenesis following cardiac ischemic episodes. Findings from recent studies support this hypothesis. In rodents, Kloth and colleagues observed a clear upregulation of *Piezo2* in stressed cardiac tissue ^98^. Further studies describe upregulation of Piezo2 in pharmacologically stressed cultured human coronary artery endothelial cells ^80^ and in cardiac samples from patients with heart failure ^99,100^.

In summary, our study elucidates the indispensable role of PIEZO2-dependent mechanosensitive signaling in coronary vasculature formation and highlights its implications for cardiac development and disease. Our study also raises a new issue for the use of therapeutics targeting PIEZO2 in the somatosensory system ^101^ as such approaches may have serious consequences for cardiovascular (patho)physiology.

## Methods

### Animal care

Experiments involving animals were performed following the German Animal Protection Act. All mice were housed in a 12:12h light/dark cycle and food/water was available *ad libitum*. To collect embryos at a determinate developmental stage, timed matings were set up with mice older than 10 weeks of age. The day of the plug was stipulated as E0.5 and pregnant females were sacrificed, at the desired embryonic stage, by cervical dislocation according to the German Animal Protection Act.

### Transgenic and mutant mice

*Piezo2*-tdTomato mice were achieved by crossing the previously described Piezo2-Cre [The Jackson laboratory, 027719 ^32^ with the Ai14 [The Jackson laboratory, 007908^102^. In this study, two Piezo2 mutant mouse models have been studied, the *Piezo2^-/-^* [The Jackson laboratory, 027720 ^32^ and the *Piezo2^R^*^275^*^6H/R2756H^* ^73^.

### Immunohistochemistry

Mice embryonic hearts were harvested and followed by fixation in 4% paraformaldehyde in PBS for 2 hours at room temperature. Tissues were washed in 1x PBS and subsequently cryopreserved in 30% sucrose. Then the hearts were frozen in OCT (Tissue-Tek®, Sakura Finetek) and sectioned on a Leica CM1950 cryostat at 10µm for immunofluorescence staining. Samples were permeabilized with 0.1% Triton X-100 for 30 minutes and blocked with 10% donkey serum (Biowest) and 1% bovine serum albumin (Sigma-Aldrich) for 1 hour at room temperature. Then, sections were incubated with primary antibodies diluted in PBST with 1% serum overnight at 4°C and with secondary antibodies for 1 hour at room temperature. DAPI (62248, Invitrogen) was used for nuclei counterstaining and DAKO fluorescent (S302380-2, Agilent) was used as the mounting medium.

E13.5 hearts whole-mount immunostainings were performed in 2 mL Eppendorf tubes in constant rotation. Hearts were firstly permeabilized with 0.5% Triton X-100 in PBS (PBST) for 1 hour and blocked with 10% Donkey serum in 0.5% PBST for 2 hours at room temperature. Samples were incubated in primary and secondary antibody cocktails for 24 hours at 4°C. In between incubations, the tissue was washed with PBST. All samples were counterstained with DAPI. Finally, the stained hearts were mounted in a µ-Slide 8 well-high glass bottom (IBIDI) using Fluoromount G (00-4959-52, eBioscience) as the mounting medium.

### Tissue clearing and immunolabelling

E18.5 *Piezo2*-tdTomato hearts were cleared following the SmartClear protocol from LifeCanvas Technologies, as it preserves endogenous fluorescence. Previously to the clearing, all PFA-fixed samples were treated with the additional epoxy-based fixation step SHIELD, as a preservation step before the delipidation (clearing) ^103^ Hearts were cleared for 3 days at 42°C using the SmartClear device (LifeCanvas Technologies). Afterwards, the samples were incubated overnight at 37°C in 50% EasyIndex + 50% distilled water with shaking, followed by a final incubation in 100% EasyIndex at 37°C overnight, for refractive index matching.

E11.5 *Piezo2*-tdTomato and E18.5 *Piezo2^-/-^* hearts were cleared and labeled following an optimized iDISCO protocol ^104^. Briefly, samples were dehydrated, washed and incubated in 66% dichloromethane/33% methanol overnight at room temperature. Afterwards, the samples were bleached overnight in 5% H_2_O_2_ in methanol, rehydrated, washed and permeabilized. Then, the hearts were blocked and incubated in the antibody solution for 4 days at 37°C. Subsequently, the samples were embedded in 1% low-melting agarose, dehydrated and incubated for 2.5 hours in 66% dichloromethane/33% methanol at room temperature. Finally, the samples were incubated in 100% dichloromethane and stored at room temperature in Ethylene Cinnamate.

### Antibodies

The following antibodies were used for immunostaining: goat anti-PECAM1 (1:250, AF3628, R&D Systems), rabbit anti-DACH1 (1:100, 10910-1-AP, Proteintech), rat anti-EMCN (1:100, sc-65495, Santacruz), rabbit anti-FABP4 (1:100, ab13979, Abcam), goat anti-SOX17 (1:100, AF1924, R&D Systems), rabbit anti-RFP (1:1000, 600-401-3979S, Rockland), rat anti-VE-Cadherin (555289, BD-Pharmigen), mouse anti-SMA (1:200, A5228, Sigma-Aldrich), mouse anti-SMA-Cy3 (1:250, C6198, Sigma-Aldrich), Alexa Fluor 488 rat (1:500, X, Abcam), Alexa Fluor 555 rabbit (1:500, ab150074, Abcam), Alexa Fluor 647 rabbit (1:500, ab150075, Abcam), Alexa Fluor 647 goat (1:500, ab150131, Abcam), Alexa Fluor 647 rat (1:500, ab150155, Abcam).

### Hematoxylin and eosin staining

Hematoxylin and eosin staining were performed, following manufacturer instructions on 10µm thick cryosections. Briefly, sections were stained with filtered 0.1% Mayers Hematoxylin for 10 minutes and then, rinsed and stained with 0.5% Eosin for 3 minutes followed by a series of washings with ddH_2_O and increasing Ethanol/ddH_2_O solutions. Finally, slides were dipped several times into RotiHistol and mounted with Roti^®^-Mount. Stained sections were imaged using a Leica DM5000B microscope.

### Single-molecule fluorescent in situ hybridization

Single-molecule fluorescent in situ hybridization was carried out according to the manufacturer’s instructions (RNAscope^TM^ Multiplex Fluorescent V2 assay, 323110, ACD) on E13.5 wild-type C57Bl/6 embryonic heart sections. *Piezo2* (Probe-Mm-*Piezo2*, 400191, ACD) and *Dach1* (Probe-Mm-*Dach1*,12071-C3, ACD) expression was studied. Opal^TM^ 570 (FP1488001KT, Akoya Biosciences) and Opal^TM^ 690 (FP1497001KT, Akoya Biosciences) fluorophores were used.

### Confocal imaging

All immunofluorescent stained tissue sections and the E11.5 and E13.5 whole-mount immunolabelled hearts were imaged using the LSM700 confocal microscope and the ZEN software. The objectives used for the imaging acquisition were the 10x NA 0.3 EC Plan-Neofluar, 20x NA 0.8 Plan Apochromat and 40x NA 1.4 Oil Plan Apochromat objectives. For all samples DAPI was excited at 405 nm (detection at 420-450 nm), Alexa Fluor 488 was excited by a 488 nm laser (detection at 500 - 550 nm), Alexa Fluor 555 was excited by a 555 nm laser (detection at 570 - 620 nm), Alexa Fluor 647 was excited by a 639 nm laser (detection at 660 - 730 nm) with a pinhole set to 1 AU.

### Light sheet microscopy

Cleared and immunolabelled whole E18.5 hearts were imaged with a Zeiss Lightsheet 7 microscope, with the ZEN 3.1 (black edition) LS software. The acquisition was done with dual side illumination (6.08 µm light sheet thickness), using LSFM 5x/0.1 foc objectives for illumination, and an EC Plan-Neofluar 5x/0.16 foc lense for detection, with the correction collars adjusted to the correct refractive index, depending on the immersion used (1.46 for EasyIndex, 1.56 for Ethylene Cinnamate). Acquisition was done sequentially, using solid state lasers 405 nm and 561 nm (camera beamsplitter LP 510 nm, emission filters BP [420-470] nm / [575-615] nm), with a PCO.Edge 5.5 sCMOS camera (6.5 µm/pixel, 1920 x 1920 pixels), and using a zoom of 0.96 (0.97 x 0.97 x 2.00 µm per voxel). Post-acquisition processing was performed with the ZEN 3.4 (blue edition) software to fuse the dual-side light sheets of each sample while the FIJI BigStitcher plug-in ^105^ was used to stitch the tiles.

### Image analysis

Identical settings for laser power, detector and pixel size were used for all samples subjected to either qualitative or quantitative analysis. For those samples where image post-processing (FIJI software) was needed, all parameters were equally modified through all the samples.

The thickness of the ventricular wall was measured from *Piezo2*^+/+^ and *Piezo2*^-/-^ E18.5 hearts longitudinally sectioned. Six sections per animal were selected, three sections containing the pulmonary artery and 3 sections with the aorta. Measurements were taken, using FIJI software, from the left ventricle, the right ventricle and the interventricular septum. A specific region of interest (ROI) below the atria was selected and the thickness of the compact and trabecular myocardium was measured. In total ten measurements were taken per area/section. The average thickness per area/animal was used in the final quantification.

The diameter of the right and left coronary arteries was measured using FIJI software from whole-mount E18.5 *Piezo2*^+/+^ and *Piezo2*^-/-^ immunolabelled hearts. Measurements were taken at the most proximal part of the coronary artery, precisely at 200 µm, 300 µm, 400 µm, 500 µm and 600 µm from the connection of the artery with the aorta. The resulting five values per artery were averaged and used for the statistical analysis.

The diameter of the pulmonary artery and the aorta were measured using FIJI from optical sections obtained from whole-mount E18.5 *Piezo2*^+/+^ and *Piezo2*^-/-^ immunolabelled hearts. Two measurements were taken per vessel and average. The resulting values were used for the statistical analysis.

Coronary vasculature surface reconstruction was performed based on the iDISCO-cleared and αSMA immunolabeled *Piezo2^+/+^* and *Piezo2^-/-^* hearts. The surface package of the IMARIS software was used to semi-automatically detect, segment, and reconstruct the immunofluorescence signal. The segmentation and 3D reconstruction were used to qualitatively study the coronary architecture.

### Echocardiography

The ultrasound examination was performed on anaesthetized 10-week-old *Piezo2^+/+^* and *Piezo2^R2756H/R2756H^* mice. Anesthesia was induced with 3-4% isoflurane (anesthesia box) and maintained with 1.5-2% isoflurane (mask ventilation). After anesthesia induction, the examination area was depilated using a shaver and hair removal cream. To prevent the eyes from drying out, an eye ointment (Bepanthen) was applied. Then, the animals were placed on a warming plate for the entire duration of the examination. Moreover, a heat lamp was used to prevent the animals from cooling down. The body temperature was monitored using a rectal probe. During the ultrasound examination, an electrocardiogram was derived and recorded to monitor the heart rate and anesthesia depth. The examination was performed on the Vevo 3100 ultrasound machine (VisualSonics Fujifilm) together with the MXD550D ultrasonic probe. Contact gel was used between the transducer and the animal. The entire examination took about 45 min, and after that, the animals woke up under supervision in a warm and quiet environment.

### Primary cardiac endothelial cells isolation

Primary cardiac endothelial cells (cEC) were obtained from E13.5 *Piezo2-*tdTomato embryos. Each isolation consisted of a pool of at least 10-15 hearts. Hearts were harvested and the atriums and outflow tract were carefully removed, resulting in the isolation of the ventricles. Subsequently, the ventricles are mechanically dissociated and the minced tissue is filtered through a 70 µm filter. The retained tissue is then enzymatically digested in Collagenase/Dispase solution (10269638001, Sigma Aldrich) for 45 minutes at 37°C, homogenized and filtered into a falcon containing 5mL of DMEM, 10% FBS (P40-37500, PAN-Biotech) and Gentamicin (G1272, Sigma Aldrich). Next, 5 mL of the same solution was added to the falcon, and centrifuge during 5’ at 1200 rpm. The supernatant was then discarded, the pellet resuspended in 1 mL of PBS-BSA 0.5% and transferred to an Eppendorf to be centrifuged for 5’ at 300g. The resulting pellet was resuspended in 100µL of PBS-BSA 0.5% and 100µL of bead/antibody solution was added and carefully homogenized. After incubating for 30 minutes at room temperature, 800µL of PBS-BSA 0.5% are added to the mix, and using the magnetic rack, the beads coupled to the cells are washed several times with MLEC medium. MLEC medium contains 400 mL DMEM-F12 (21041-025, Invitrogen), 20% FBS inactivated, 1% penicillin/streptomycin (15140122, Gibco) and 4 mL ECGS-H (C-30120, Promocell). Finally, the cells are resuspended in 500 µL of MLEC and each pool is seeded in one well of a 24-well plate coated with 0.5% gelatin (G1393, Sigma Aldrich). The next day, cells are washed with 1 mL of PBS-BSA 0.5% and 500 µL of MLEC is added to the well. Once cells reached confluency, usually after one week in culture, they were dissociated with trypsin and seeded on the pillar arrays for the electrophysiology experiments.

To prepare the bead/antibody solution, 6µL of sheep anti-rat IgG Dynabeads (11035, Dynal Invitrogen) per pool of hearts will be placed in 1 mL of PBS-BSA 0.5%, washed several times and resuspended in the same volume of PBS-BSA 0.5% as beads (6µL/pool). Then, 1.5 µL/pool of VE-Cadherin antibody is added to the beads solution and incubated for 1 hour at room temperature with soft agitation. After incubation, 100 µL of PBS-BSA 0.5% per pool was added and softly homogenized.

### Preparation of pillar arrays

Pillar arrays were prepared following the established protocol ^63^. In summary, silanized negative masters served as templates, which were subsequently coated with polydimethylsiloxane (PDMS) from the syligard 184 silicone elastomer kit (Dow Corning Corporation) mixed with a curing agent in a 10:1 ratio (elastomeric base:curing agent). The mixture was incubated for 30 min, and glass coverslips were positioned on the top of negative masters containing PDMS, followed by banking for 1h at 110°C. Subsequently, pillar arrays were peeled from the negative masters. Each pilus within the array exhibited a radius of 1.79 µm and a length of 5.8 µm. Prior the cell culture use, the pillar arrays underwent plasma cleaning using a Femto low-pressure plasma system (Deiner Electronic GmbH) and were salinized using vapor phase (tridecafluoro-1,1,2,2-tetrahydrooctyl trichlorosilane 97% (AB111444, ABCR GmbH & Co. KG, Karlsruhe, Germany) for 45 min, followed by a coating with laminin EHS laminin and Poly-L-lysine in a 1:1 ratio (v:v) for at least 2h at 37°C.

### Pillar arrays experiments

Whole-cell patch clamp recordings were conducted on yolk sac (YS) and cEC cells (isolated as described above) using borosilicate glass pipettes (Harvard apparatus, 1.17 mm x 0.87 mm) with a resistance of 3-6 MΩ). The pipettes were filled with the intracellular solution containing (in mM): 110 KCl, 10 NaCl, 1 MgCl_2_, 1 EGTA and 10 HEPES; the pH was adjusted to 7.3 with KOH. The extracellular solution contained (in mM): 140 NaCl, 4 KCl, 2 CaCl_2_, 1 MgCl_2_, 4 Glucose and 10 HEPES; pH adjusted to 7.4 with NaOH. Pipette and membrane capacitances were compensated using the auto-function of Patchmaster (HEKA, Elektronik GmbH, Germany) while series resistance was compensated to minimize voltage errors. For pillar arrays, currents were recorded at a holding potential of -60 mV. Currents were recorded at 10 KHz and filtered at 3 KHz using an EPC-10 USB amplifier (HEKA, Elektronik GmbH, Germany) and Patchmaster software.

A single pilus was deflected using a heat-polished borosilicate glass pipette driven by an MM3A micromanipulator (Kleindiek Nanotechnik, Germany) as previously described in Poole et al., 2014 ^63^. Pillar deflection stimuli range from 1-1000 nm, with larger deflections discarded. For quantification and comparison analysis, the data were binned by the magnitude of the stimuli (1-10, 11-50, 51-100, 101-250, 251-500, 501-1000 nm), and the mean current amplitudes within each bin were calculated for every cell. Deflection-gated currents were classified according to their inactivation kinetics: rapidly adapting (RA, τinact < 5 ms), intermediate adapting (IA, τinact 5-50 ms), and slowly adapting currents (SA, τinact > 50 ms).

To calculate pillar deflection, bright field images (Zeiss 200 inverted microscope) were captured using a 40X objective and CoolSnapEZ camera (Photometrics, Tucson, AZ) before and after every pillar stimulus. Pillar deflection was determined by comparing the light intensity of the center of each pilus before and after every stimulus using a 2D-Gaussina fit (Igor Software, WaveMetrics, USA).

High-Speed Pressure Clamp (HSPC) experiments. HSPC (Ala Scientific) was conducted on excised outside-out patched from YS cells. Recording pipettes with a final resistance of 6-8 M were used. Positive pressure pulses were delivered through the recording pipette. The pressure steps protocol involved a series of stimuli ranging from 10 to 150 mmHg, in 20 mmHg increments, while maintaining the patch potential at -60 mV. Recording solutions were prepared in symmetrical ionic conditions containing (in mM): 140 NaCl, 10 HEPES, 5 EGTA adjusted to pH 7.4 with NaOH.

For both methods, currents and the biophysical parameters were analyzed using FitMaster (HEKA, Elektronik GmbH, Germany).

### Single-cell RNA sequencing

Gene count matrices from single-cell sequencing from E12, E15, P2 and 8-weeks-old C57Bl/6J animals were generated by Cano et al. ^48^, and deposited on the Gene Expression Omnibus repository with accession number GSE223266. The sequence data were mapped to the mouse reference genome (mm10 pre-build references v 2.1.0). Custom code was generated using R to analyze the data and to generate plots as described in Cano et al. ^48^.

### Statistical analysis

All data analyses were performed using GraphPad Prism and tested for normality. For normally distributed data, statistically significant differences were determined with the parametric, unpaired Student’s t-test, whereas not normally distributed data were compared using the non-parametric Mann-Whitney-test. For all cases, the standard error of the mean (SEM) is provided. The term significant was used if the p-value was less than 0.05 (p<0.05). Exact p-values and N values are reported in the figure legends.

## Acknowledgements

We thank Anke Scheer and Franziska Bartelt for their help with mouse genotyping. We thank Stephanie Rode, Franziska Westphal and Franziska Kratz for animal husbandry. We thank Michael Gotthardt and members of the Hammes and Lewin labs for constructive comments on the manuscript.

## Author contributions

Conceptualization: GRL and AH

Mouse model design: AH, GRL, MPP, and OSC

Developmental characterization, immunohistochemistry, light sheet imaging and smFISH: MPP, CF, EC, SR, AS

Endothelial cell patch clamp physiology: OSC and MPP

In vivo cardiac ultrasound: MT and AH

Single cell sequencing and analysis: EC, and ELL.

Supervision: AH, GRL, NH, HG, and AH

Funding acquisition: AH, GRL, NH and HG

Writing AH, MPP and GRL with input from all authors

## Data and materials availability

All data are available in the main text or the supplementary materials.

## Extended data

**Fig. 1.**
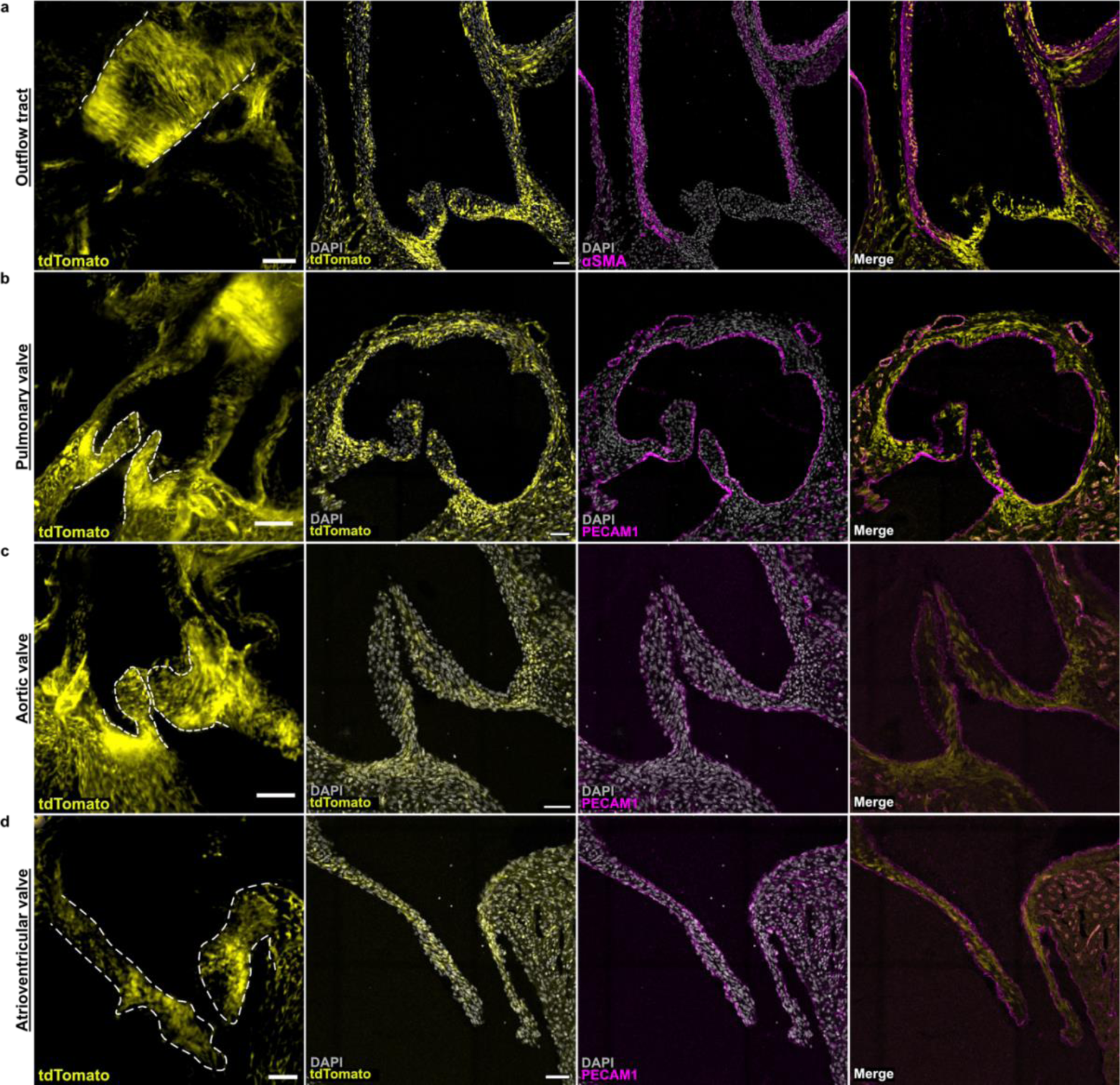
*Piezo2*-driven tdTomato is expressed in the outflow tract and the cardiac valves. **a - d,** Outflow tract and cardiac valves three-dimensional rendering and immunolabelled sections from an E18.5 tdTomato^+^ heart. tdTomato signal (yellow) in the outflow tract (**a**) colocalizes with αSMA (magenta). In the cardiac valves (**b - d**), tdTomato+ cells (yellow) are surrounded by endothelial cells (PECAM1 positive, magenta), indicating that they are likely valve interstitial cells (n = 2, Scale bar: 100µm [3D] and 50µm [sections]).

**Fig. 2.**
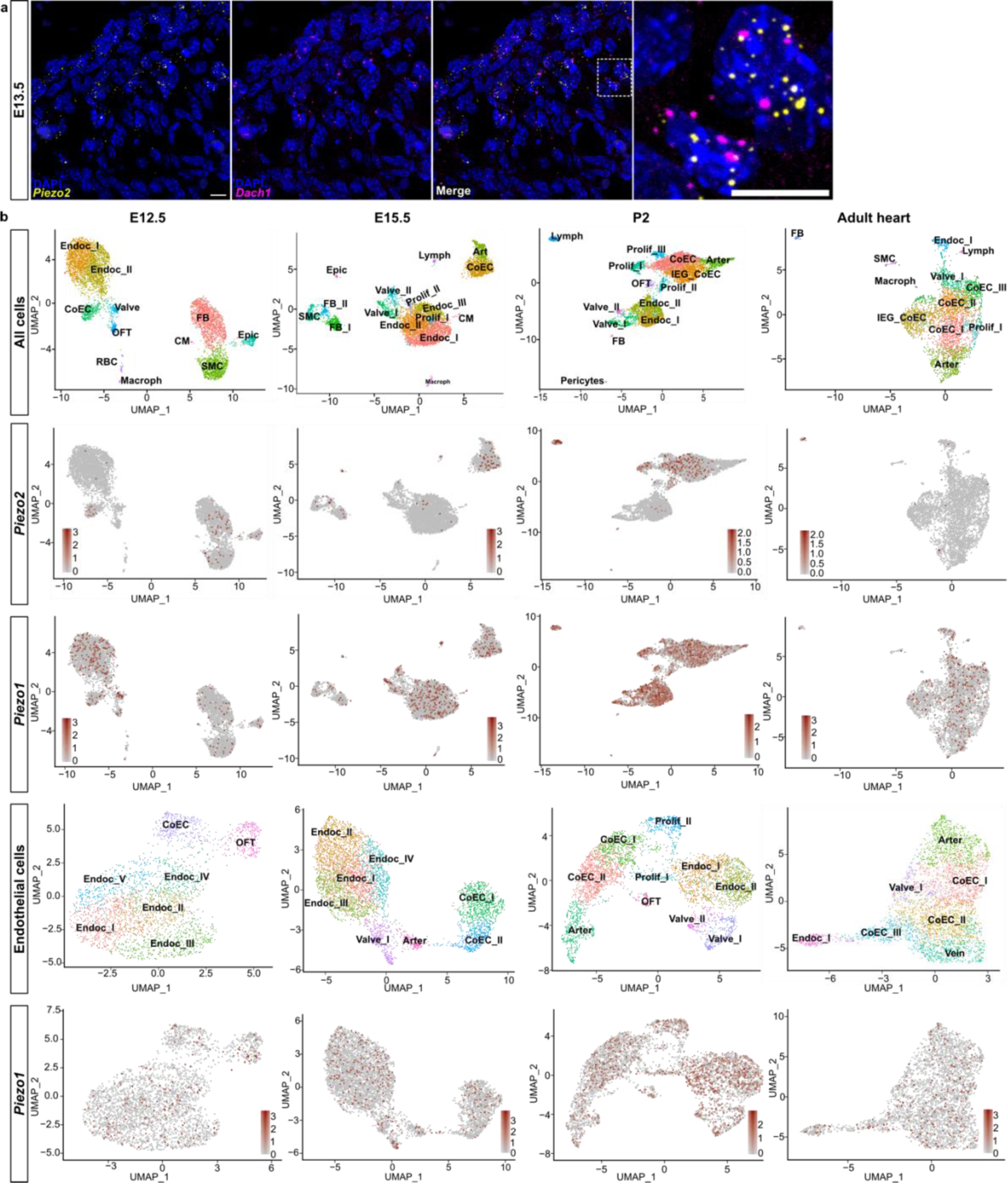
a, Expression of *Piezo2* mechanosensitive ion channels in heart tissue. Single-molecule in situ hybridization showing coronary endothelial cells expressing *Piezo2* and *Dach1* at E13.5. Nuclei are labeled with DAPI. Scale bar: 10 µm. **b**, **scRNAseq identifies distinct cluster representations for *Piezo2^+^* and *Piezo1^+^*cardiac cells.** UMAP of cardiac endothelial cells from E12, E15, P2 and murine adult hearts (8 weeks old) showing the following clusters: endocardium (Endoc_I to Endoc_V), coronary endothelium (CoEC_I to CoEC III), proliferating cells (Prolif_I to Prolif_II), valvular endocardium (Valve_I and Valve_II), arterial cells (Arter), venous cells (Vein), and outflow tract cells (OFT). Cell clusters: endocardium (Endoc_I to Endoc_V), coronary endothelium (CoEC_I to CoEC III), proliferating cells (Prolif_I to Prolif_II), valvular endocardium (Valve_I and Valve_II), arterial cells (Arter), venous cells (Vein), and outflow tract cells (OFT).

**Fig. 3.**
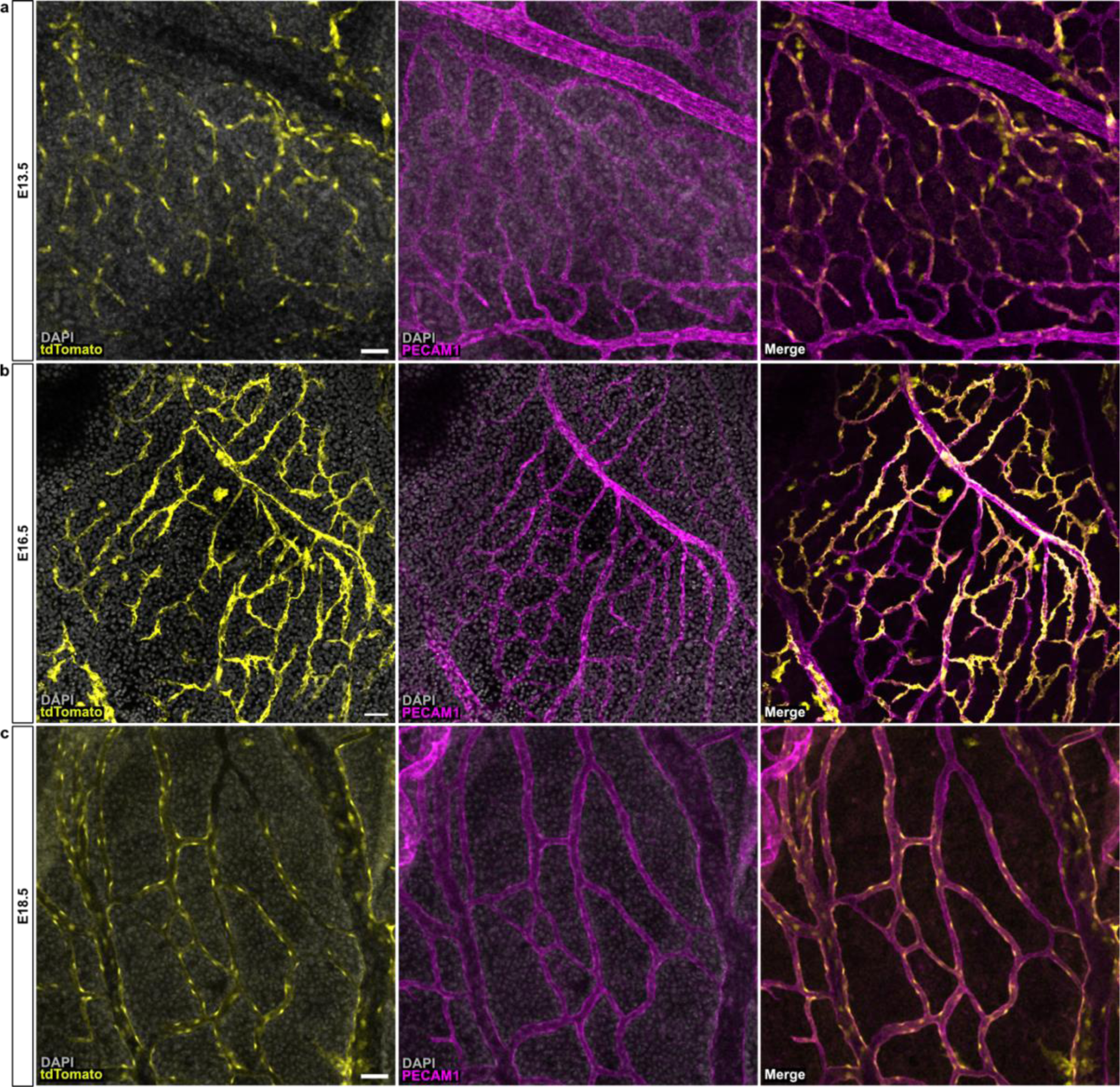
Expression of Piezo2-driven tdTomato in the developing yolk sac. **a - c**, Immunolabeling of whole yolk sacs showing tdTomato signal (in yellow) colocalized with the endothelial marker PECAM1 (in magenta) at embryonic days E13.5 (**a**), E16.5 (**b**), and E18.5 (**c**) (n = 4 per stage, Scale bar: 50 µm).

**Fig. 4.**
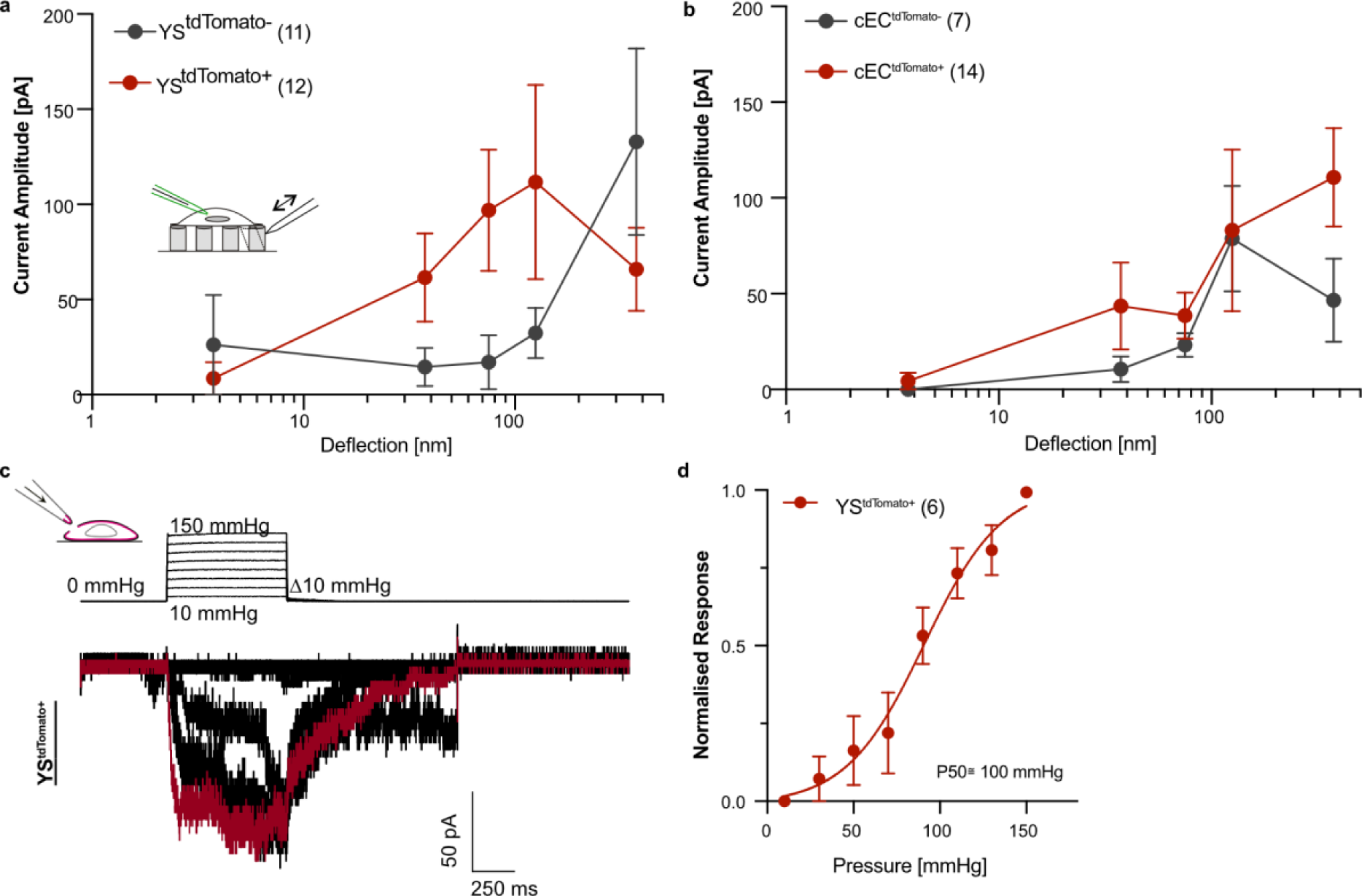
Mechanosensitive currents measured in YS^tdTomato+^ and cEC^tdTomato+^ cells. **a - b**, Deflection-current amplitude relationships of the yolk sac (YS) cells (**a**) and cardiac endothelial cells (cEC). **b**, Graph showing that deflection gated currents from tdTomato^+^ and tdTomato^-^ cells displayed similar mechanical responses (Two-way ANOVA, P > 0.05). **c**, Representative trace of stretch-sensitive currents stimulated with a pressure-step protocol from outside-out patches in YS^tdTomato+^ cells at a resting membrane potential of -60 mV. **d**, Stretch-response curve of the endogenous stretch sensitive currents in YS^tdtomato+^ cells. Peak currents were normalized to their maximum.

**Fig. 5.**
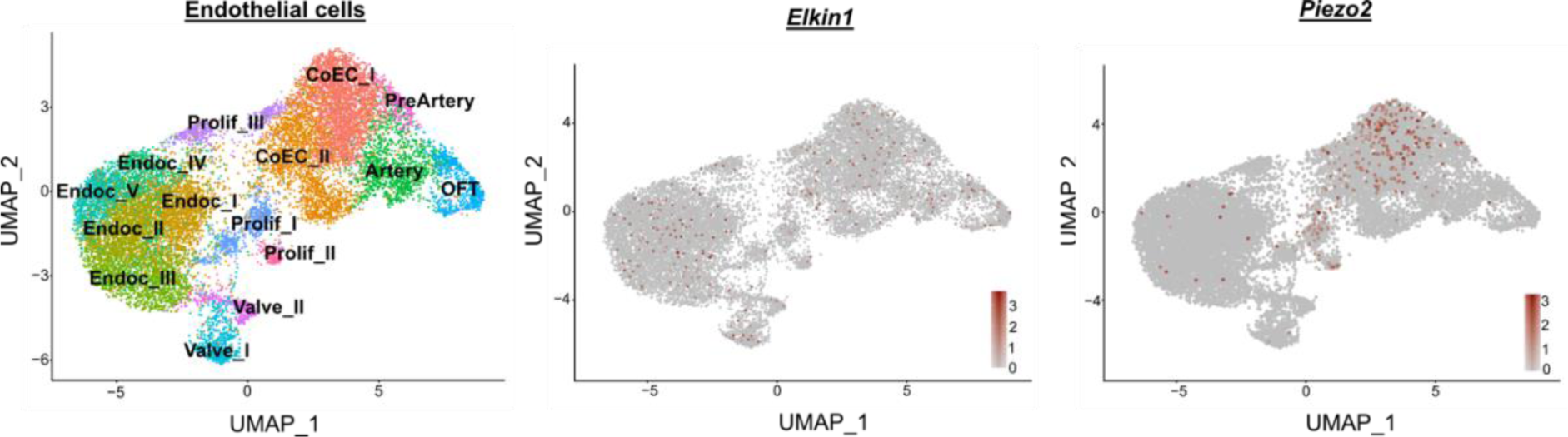
Expression of *Elkin1*. scRNAseq of E12, E15, P2 and adult hearts identify *Elkin1* expression in endothelial cell clusters that are *Piezo2^+^.* Cell clusters: endocardium (Endoc_I to Endoc_V), coronary endothelium (CoEC_I to CoEC III), proliferating cells (Prolif_I to Prolif_II), valvular endocardium (Valve_I and Valve_II), arterial cells (Arter), venous cells (Vein), and outflow tract cells (OFT).

**Fig. 6.**
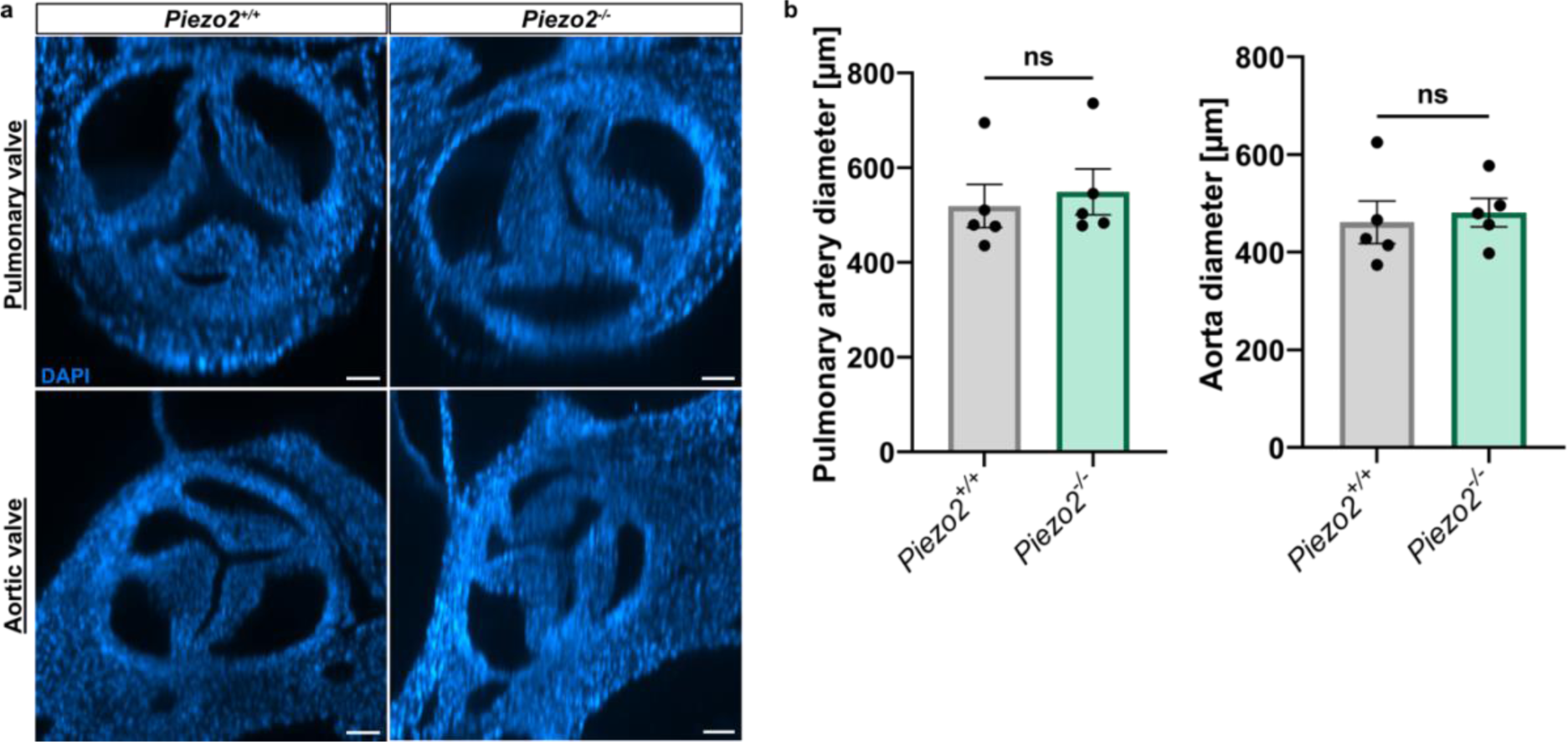
*Piezo2* mutations do not alter the cardiac valve and OFT morphology. **a**, Representative images of the aortic and pulmonary valves of *Piezo2^+/+^* and *Piezo2^-/-^*E18.5 embryos. Optical sections from whole mount light sheet images are shown. Nuclei are stained with DAPI **b**, *Piezo2*^-/-^ mutants present a similar pulmonary artery and aorta diameter compared to Piezo2^+/+^ (n = 5 per genotype, Mann-Whitney test, p = 0.420 [Pulmonary artery] and p = 0.547 [Aorta]. Scale bars: 100µm).

**Fig. 7.**
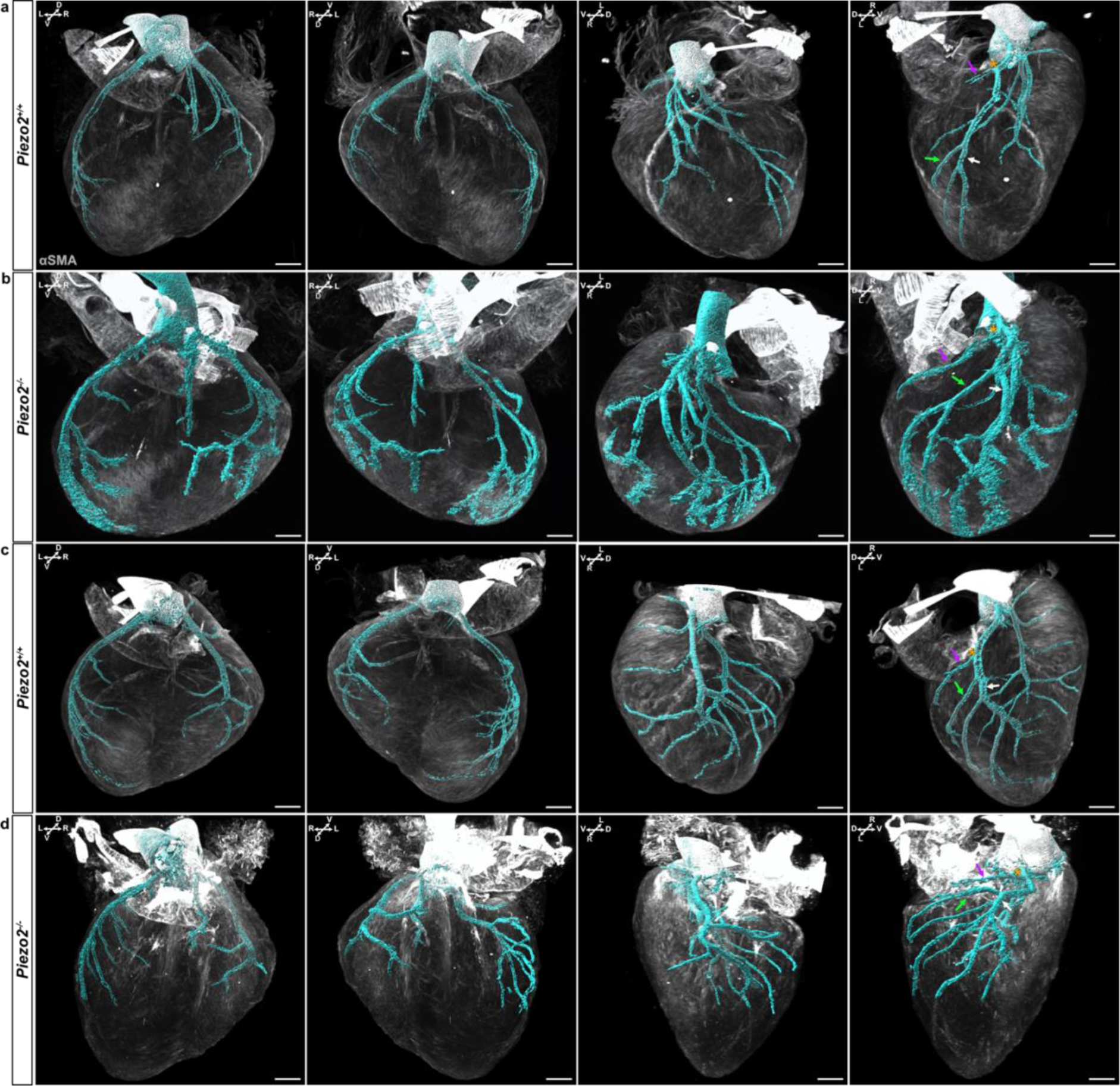
Three-dimensional rendering of *Piezo2^+/+^* and *Piezo2^-/-^* E18.5 hearts. **a - d,** iDisco-cleared and αSMA (gray) immunolabelled hearts from *Piezo2^+/+^* wild types (**a** and **c**) and *Piezo2^-/-^* mutants, used for coronary vasculature reconstruction (cyan) (Scale bars: 250µm). Arrows indicate the left coronary artery branches: the circumflex artery (magenta arrow), the diagonal branch (green arrow) and the left descending artery (white arrow). The orange asterisk shows the branching point of the circumflex artery.

## Extended Data

### Movies

**Movie 1**. **Piezo2-driven fate mapping defines the coronary vasculature.** Video recording (360° horizontal and 360° vertical views) showing the coronary vasculature of a representative tdTomato^+^ (yellow) E18.5 heart.

**Movie S2. Piezo2-driven fate mapped coronary vessels connect to the aorta.** Video recording of the optical view of a representative whole-mount tdTomato^+^ E18.5 heart.

**Movies 3 to 8. Three-dimensional rendering of *Piezo2^+/+^* and *Piezo2^-/-^* E18.5 hearts**. Video recording (360° horizontal view) of three Piezo2^+/+^ hearts (S3 to S4) and three Piezo2^-/-^ hearts (S5 to S8) with the coronary vasculature reconstructed.

### Supplementary data

Source data

### Tables

**Table 1.** E12.5 Single cell read data used Fig. 3c

**Table 2.** E15.5 Single cell read data used Fig. 3c

**Table 3.** P2 Single cell read data used Fig. 3c

